# Antibody targeting of aging damaged isoDGR-proteins doubles lifespan in a mouse model of chronic inflammation

**DOI:** 10.1101/2023.03.11.532237

**Authors:** Pazhanichamy Kalailingam, Khalilatul-Hanisah Mohd-Kahliab, SoFong Cam Ngan, Ranjith Iyappan, Evelin Melekh, Tian Lu, Gan Wei Zien, Bhargy Sharma, Tiannan Guo, Adam J. MacNeil, Rebecca E. K. Macpherson, Evangelia Litsa Tsiani, Deborah D. O’Leary, Kah Leong Lim, I Hsin Su, Yong-Gui Gao, A Mark Richard, Raj N. Kalaria, Christopher P. Chen, Neil E. McCarthy, Siu Kwan Sze

**Affiliations:** School of Biological Sciences, Nanyang Technological University, Singapore; Department of Health Sciences, Faculty of Applied Health Sciences, Brock University, St. Catharines, ON, L2S 3A1, Canada; iMarker lab, Westlake Laboratory of Life Sciences and Biomedicine, Key Laboratory of Structural Biology of Zhejiang Province, School of Life Sciences, Westlake University, Hangzhou, Zhejiang Province, China; Lee Kong Chian School of Medicine, Nanyang Technological University, Singapore; Department of Cardiology, National University Heart Centre, Singapore, Singapore; Department of Cardiology, University of Otago, Christchurch, New Zealand; Institute of Neuroscience, Campus for Ageing and Vitality, Newcastle University, Newcastle upon Tyne, United Kingdom, NE4 5PL; Memory, Aging and Cognition Centre, National University Health System, Singapore; Centre for Immunobiology, The Blizard Institute, Bart’s and The London School of Medicine and Dentistry, Queen Mary University of London, United Kingdom, E1 2AT

**Keywords:** immunotherapy, isoDGR, Pcmt1, inflammation, lifespan, antibody, immune clearance, ADCP, anti-isoDGR

## Abstract

Aging is the result of the accumulation of molecular damages that impair normal biochemical activities. We previously reported that aging-damaged amino acid sequence NGR (Asn-Gly-Arg) results in a ‘gain-of-function’ conformational switching to isoDGR (isoAsp-Gly-Arg) motif. This integrin-binding motif activates leukocytes to induce chronic inflammation, which are characteristic features of age-linked cardiovascular disorders. We now report that anti-isoDGR immunotherapy doubles lifespan in mouse model of chronic inflammation. We observed extensive accumulation of isoDGR and inflammatory cytokine expression in multiple tissues from Pcmt1-KO and old WT animals, which could also be induced via injection of isoDGR-modified plasma proteins or synthetic peptides into young WT animals. However, weekly injection of anti-isoDGR mAb (1mg/kg) was sufficient to significantly reduce isoDGR-modified proteins and pro-inflammatory cytokine expression, improve behaviour and coordination, and double the average lifespan of Pcmt1-KO mice. Mechanistically, isoDGR-mAb mediated the immune clearance of damaged isoDGR-proteins by antibody-dependent cellular phagocytosis. These results indicate that immunotherapy targeting aging-damaged proteins may represent effective interventions for a range of age-linked degenerative disorders.

**Graphical Abstract:** Anti-isoDGR immunotherapy induces immune clearance of aging damaged isoDGR-proteins to reduce chronic inflammation, improve behaviour and coordination, and double lifespan in PCMT^-/-^ mice.

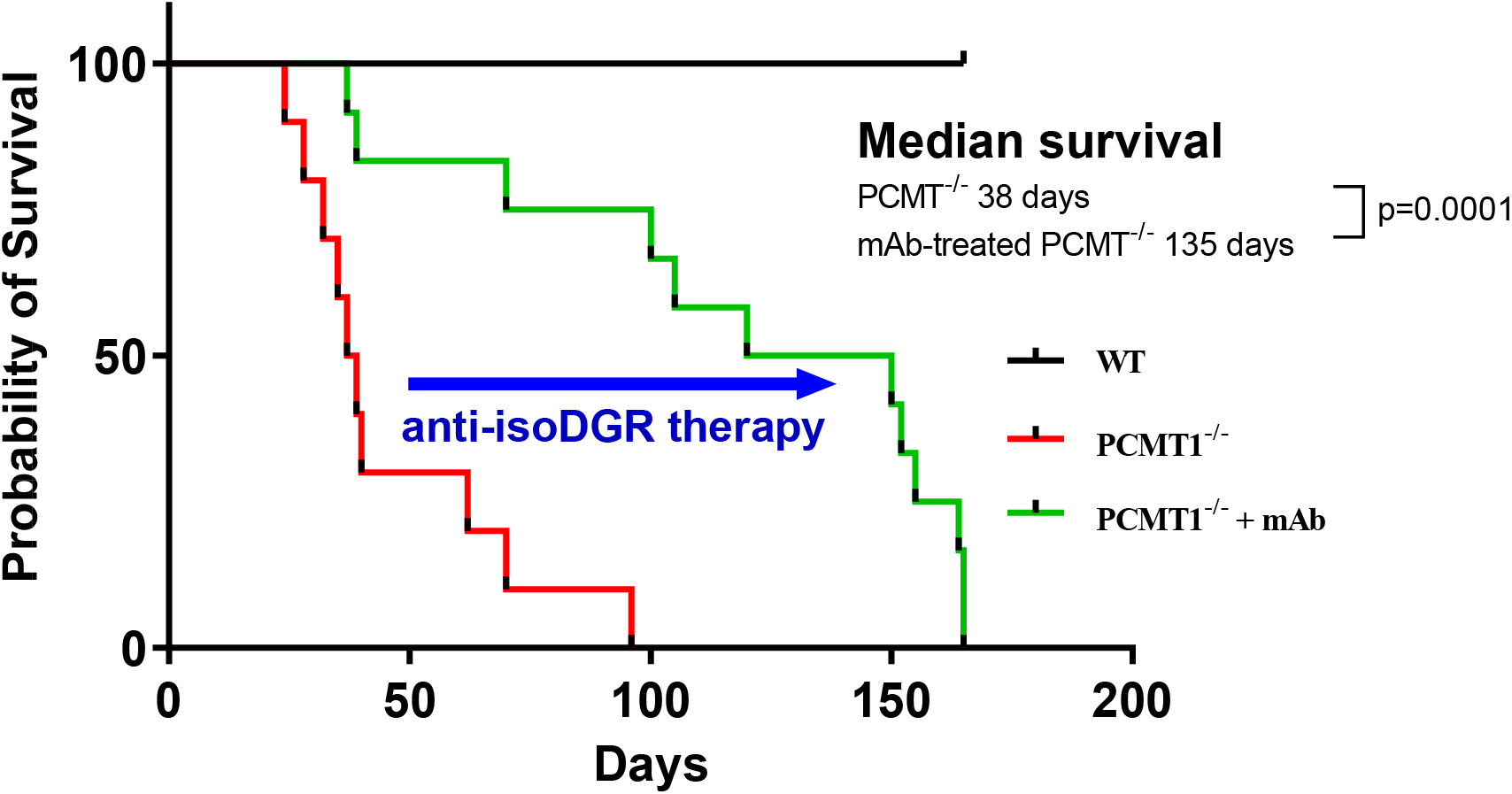

## INTRODUCTION

Aging is a complex process of time-dependent decline in key biological functions, resulting in increased susceptibility to chronic diseases and reduced lifespan. At molecular level, aging is thought to be underpinned by progressive biomolecular damage caused by protein degenerative modifications (DPMs), including oxidation, deamidation, glycation, and a range of other non-enzymatic modifications.^1–7^ We now recognize that aging is a consequence of deleterious chemical processes that damage biomolecules and impair the homeostatic functions programmed by our genomes.^8–12^ However, it is unclear whether therapeutic targeting of these damaged biomolecules represents an effective strategy for maintaining tissue function and extending healthy lifespan.

The functional impact of DPMs depends on the mode of modification and the target molecule involved. For example, deamidation leads to the accumulation of isoaspartate residues that progressively disrupt protein integrity and alter biological activity.^11,13–22^ However, ‘gain of function’ structural changes caused by DPMs may play equally important roles in human pathology.^13,23–27^ Accumulation of isoaspartate residues can occur via deamidation of asparagine or isomerization of aspartic acid residues under the influence of microenvironmental stresses, flanking amino acid sequences, and genetic factors.^11,15,20,28–31^ Consequently, DPMs greatly increase the diversity of biomolecules present in body tissues,^32^ with a high probability of generating proteoforms capable of interacting with or binding to key biomolecules in novel ways. Indeed, we recently reported that deamidation of the amino acid sequence NGR (Asn-Gly-Arg) in extracellular matrix (ECM) proteins results in ‘gain-of-function’ conformational switching to isoDGR (isoAsp-Gly-Arg) motifs^13,23,24^ that can bind to integrins and promote immune cells activation.^24–27^ In contrast to isoaspartate-modified proteins within cells that can be repaired by enzyme, long-lived extracellular matrix (ECM) proteins cannot be repaired by intracellular mechanisms and are susceptible to progressive accumulation over time.^5,7,11,15^ Accordingly, age-linked isoDGR modifications have previously been detected in fibronectin, laminin, tenascin C, and several other ECM proteins derived from human carotid plaque tissues,^23,24,33^ suggesting that these molecules may be capable of enhancing leukocyte binding to the atherosclerotic matrix.^13,24^ While age-associated DPMs have now been clearly implicated in a range of chronic diseases linked with human aging,^10^ the potential benefits of targeting these structures with specific immunotherapies remain largely unknown.

The protein l-isoaspartate (d-aspartate)-*O*-methyltransferase (Pcmt1) enzyme is expressed in all mammalian tissues and mediates the repair of age-linked protein damage (by promoting conversion of abnormal aspartyl residues to l-aspartyl forms).^14,15^ Previous studies have shown that global deletion of Pcmt1 in mice leads to the accumulation of isoaspartate in all body tissues and death at around 42 days.^14,16,34^ This models the decline in protein repair function as animals age. Having previously observed that long-lived matrix proteins undergo age-dependent deamidation,^23,24^ we hypothesized that isoDGR modification of ECM components may represent the mechanistic link in common pathologies affecting both elderly humans and Pcmt1-KO mice. We previously observed that isoDGR motifs accumulate in blood plasma and body tissues which active CD68+ macrophage to express pro-inflammatory cytokines.^13^ We also confirmed that the interaction of isoDGR-modified proteins with macrophage integrins underpinned this pathology,^24^ suggesting a potential role for this axis in the ‘inflammaging’ characteristics of advancing age.^35–37^ In the current study, we therefore tested whether anti-isoDGR immunotherapy can reduce low-grade inflammation and extend the lifespan of Pcmt1-KO mice.

Here we report that weekly injection of 1mg/kg isoDGR-specific monoclonal antibody (mAb) significantly increased body weight, improved behaviour and coordination, and doubled the average lifespan of Pcmt1^-/-^ mice. In addition, mAb treatment decreased levels of circulating pro-inflammatory cytokines and reduced tissue inflammation, strongly suggesting that isoDGR-modified proteins are at least partly responsible for the pathology observed in Pcmt1-deficient animals. We then explore the mechanisms underlying the observed phenotype changes, further *in vitro* and *in vivo* assays demonstrated that anti-isoDGR mAb can induce immune clearance of the target motif. Western blot analysis showed that anti-isoDGR mAb treatment reduces isoDGR-modified protein levels in both brain and liver tissues, indicating antibody-dependent effector functions mediated immune clearance *in-vivo*. Effector functions mediated by FcγRs include antibody-dependent cellular toxicity (ADCC) and antibody-dependent cellular phagocytosis (ADCP) which exert a critical influence on the therapeutic efficacy of antibody drugs.^38^ FACS and immunofluorescent imaging further demonstrated that the mAb-mediated immune clearance via antibody-dependent cellular phagocytosis (ADCP). Since damaged proteins are known to accumulate in body tissues with advancing age, immunotherapy targeting the damaged proteins as demonstrated here may be an effective intervention for human age-linked disorders and in extending healthy lifespans.

## RESULTS

### Anti-isoDGR immunotherapy doubles the lifespan of mouse with isoaspartate accumulation

Global deletion of repair enzyme Pcmt1 leads to tissue accumulation of isoaspartate residues and premature death of Pcmt1^-/-^ mice^14,16^, but the mechanistic basis of this pathology is not fully understood. We have previously shown that isoDGR-modified fibronectin is a critical mediator of vascular inflammation in Pcmt1^+/-^ mice via interaction with resident macrophages in an atherosclerotic CVD model.^13^ We observed that isoDGR-modified proteins positively correlated with CD68+ macrophage cells in both intima and adventitial layers of the aorta, with isoDGR engagement of macrophage integrins playing a key role in triggering inflammatory cytokine release and progression of atherosclerotic lesions.^24^ To fully elucidate the role of isoDGR motifs in age-linked disease, we generated global Pcmt1^-/-^ mice by crossing Pcmt1^+/-^ parents (25% of pups born were Pcmt1^-/-^ consistent with the expected Mendelian ratio). Genotypes were confirmed by PCR of tail genomic DNA (data not shown) and western blot analysis of liver protein extracts (Fig.1A). Pcmt1^-/-^ neonates were viable but displayed significantly reduced body weights and died prematurely compared to PCMT1^+/+^ mice. We hypothesized that isoDGR-induced tissue inflammation is partly responsible for the pathology observed in Pcmt1^-/-^ mice. We therefore tested if anti-isoDGR immunotherapy using specific mAb could be used to reduce levels of damaged proteins in body tissues from Pcmt1^-/-^ pups. Strikingly, intraperitoneal injection of 1mg/kg/week isoDGR mAb significantly increased the body weight of Pcmt1^-/-^ mice (Fig.1B-D) and doubled the average lifespan of both male and female Pcmt1^-/-^ mice compared with untreated Pcmt1^-/-^ mice (Fig.1E). These data strongly suggested that isoDGR-modified proteins can be targeted with specific immunotherapy to reduce the tissue damage and extend lifespan.

**Figure 1:**
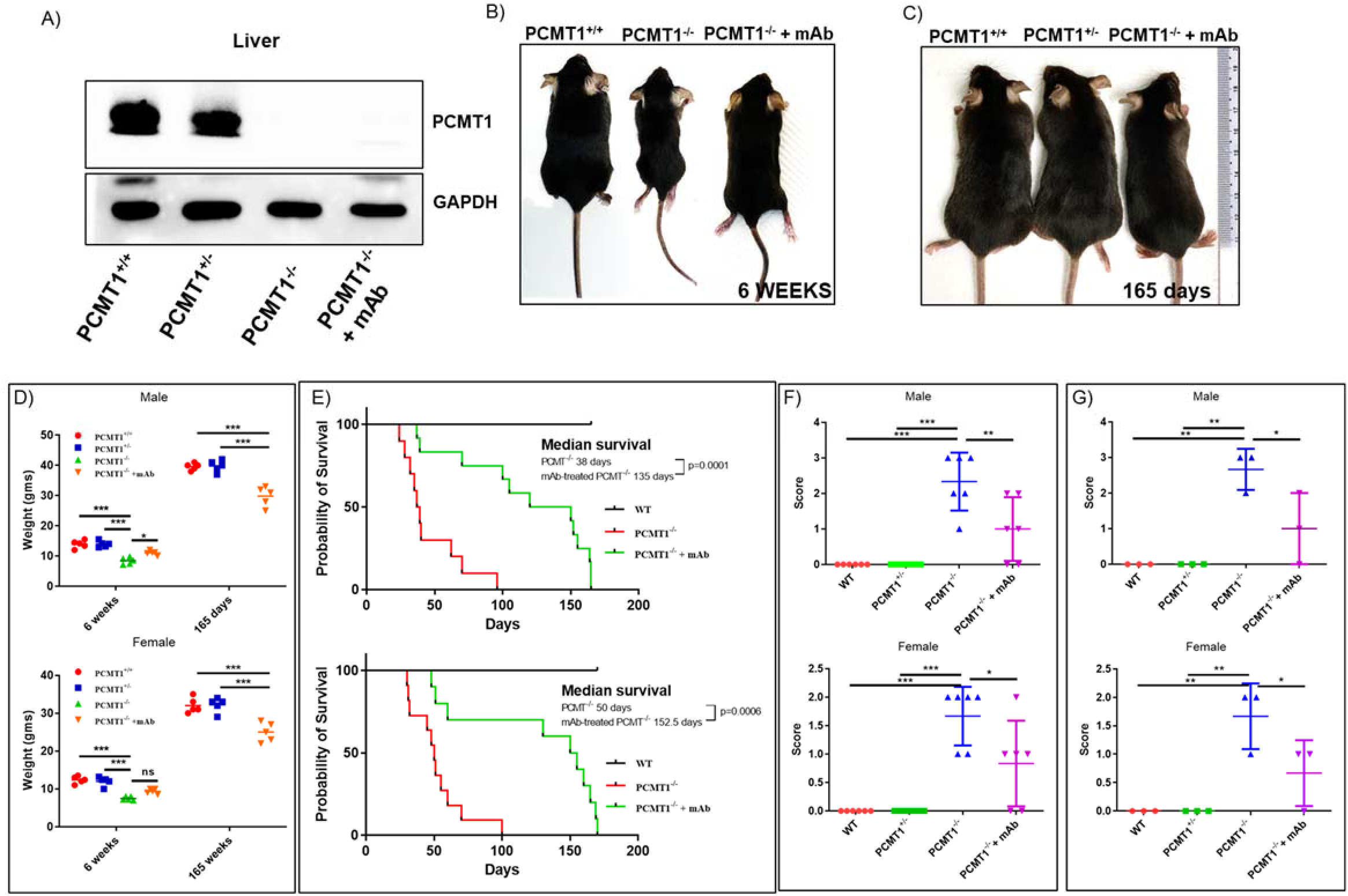
IsoDGR mAb treatment improves motor function and extends lifespan of Pcmt1^-/-^ mice. **(A)** Protein lysate from liver of Pcmt1^+/+^ (left), Pcmt1^+/-^, Pcmt1^-/-^, and mAb-treated Pcmt1^-/-^ mice (right) were subjected to western blot using antibodies against Pcmt1 (GAPDH was used as a loading control). **(B)** Representative images of Pcmt1^+/+^ (left), Pcmt1^-/-^, and mAb-treated Pcmt1^-/-^ mice (right) at 6 weeks. **(C)** Pcmt1^+/+^ (left), Pcmt1^+/-^, and mAb-treated Pcmt1^-/-^ mice (right) at 165 days of age. **(D)** Dot plot shows average body weight (male or female) for each genotype at 6 weeks and 165 days of age (n=5). **(E**) Graph shows percent survival after birth for Pcmt1^+/+^, Pcmt1^-/-^ and mAb-treated Pcmt1^-/-^ mice (male or female), Statistical differences in survival were evaluated using Mantel-Cox log-rank test. **(F)** Dot plot shows quantitative analysis of hind-limb clasping test (n=6) and **(G)** ledge scores (n=3); Pcmt1^-/-^ mice achieved higher scores than Pcmt1^+/+^ mice, while anti-isoDGR mAb treatment reduced scores in Pcmt1^-/-^ animals at 6 weeks. Statistical significances were determined by 1-way ANOVA for body weight and by Kruskal-Wallis test for hind-limb clasping and ledge experiments. Results are shown as mean ± SD (***p < 0.001, **p < 0.01, *p < 0.05).

### IsoDGR-specific antibody therapy improves motor function of Pcmt1^-/-^ mice

Deletion of Pcmt1 in mice causes severe motor dysfunction associated with seizure activity. Cytosolic proteins from all Pcmt1-deficient tissues contained significantly increased levels of protein damage compared to PCMT1^+/+^ tissues (as evidenced by 4-to 8-fold higher amounts of isoaspartyl residues).^34^ Intriguingly, protein damage accumulated to particularly high levels in brain cytosol fractions from Pcmt1^-/-^ mice.^14,39^ We therefore proceeded to test the ability of isoDGR-specific immunotherapy to preserve motor and cognitive function in Pcmt1^-/-^ mice using clasping and ledge behaviour tests.^40–42^ Six-week-old PCMT1^+/+^ and Pcmt1^+/-^ mice displayed normal extension reflex in the hind-limbs and used body torsion when suspended in the air. In contrast, Pcmt1^-/-^ mice exhibited severe hind-limb clasping and high dysfunction scores than were improved upon treatment with isoDGR-specific mAb (Fig.1F). Ledge tests also revealed impaired motor coordination in the Pcmt1^-/-^ mice, as evidenced by difficulty in paw placement and multiple limb slips during forward movement. Consequently, KO mice took longer to traverse the ledge than PCMT1^+/+^ and Pcmt1^+/-^ mice. However, administration of isoDGR-specific mAb was sufficient to improve paw placement and reduced limb slip frequency during forward movement of Pcmt1^-/-^ mice (Fig.1G). Together, these results indicate that anti-isoDGR immunotherapy improved motor function in Pcmt1^-/-^ mice.

### Anti-isoDGR mAb treatment reduces isoDGR-modified protein levels in tissues

To understand the mechanism underlying the observed phenotype changes, we next proceeded to test whether the therapeutic effects of anti-isoDGR antibody could be attributed to antigen neutralization^43^ or antibody-dependent effector functions mediated via leukocyte Fc receptors.^44,45^ If anti-isoDGR mAb simply neutralized the target motif *in vivo*, the overall abundance of isoDGR-damaged proteins in tissues should remain unchanged. Alternatively, if the beneficial effects of treatment were mediated by antibody-induced leukocyte clearance of isoDGR-damaged proteins, motif levels should be reduced in body tissues. We therefore assessed isoDGR-modified protein levels by western blot analysis of whole brain and liver lysates from 6-week-old Pcmt1-KO that had been treated or not with anti-isoDGR mAb. This analysis indicated that Pcmt1-KO mice accumulated large quantities of isoDGR-damaged proteins in both the liver and brain relative to age-matched PCMT1^+/+^ and Pcmt1^+/-^ mice (Fig.2A and 2B), but motif levels were significantly reduced by specific mAb treatment (Fig.2C and 2D). While the blood-brain-barrier can limit antibody penetration into brain tissue, we nonetheless observed a substantial reduction of isoDGR-modified proteins in mice treated with target-specific mAb (Fig.2C). In addition, anti-isoDGR immunotherapy significantly induced immune clearance of this damage motif from the liver (Fig.2D).

**Figure 2:**
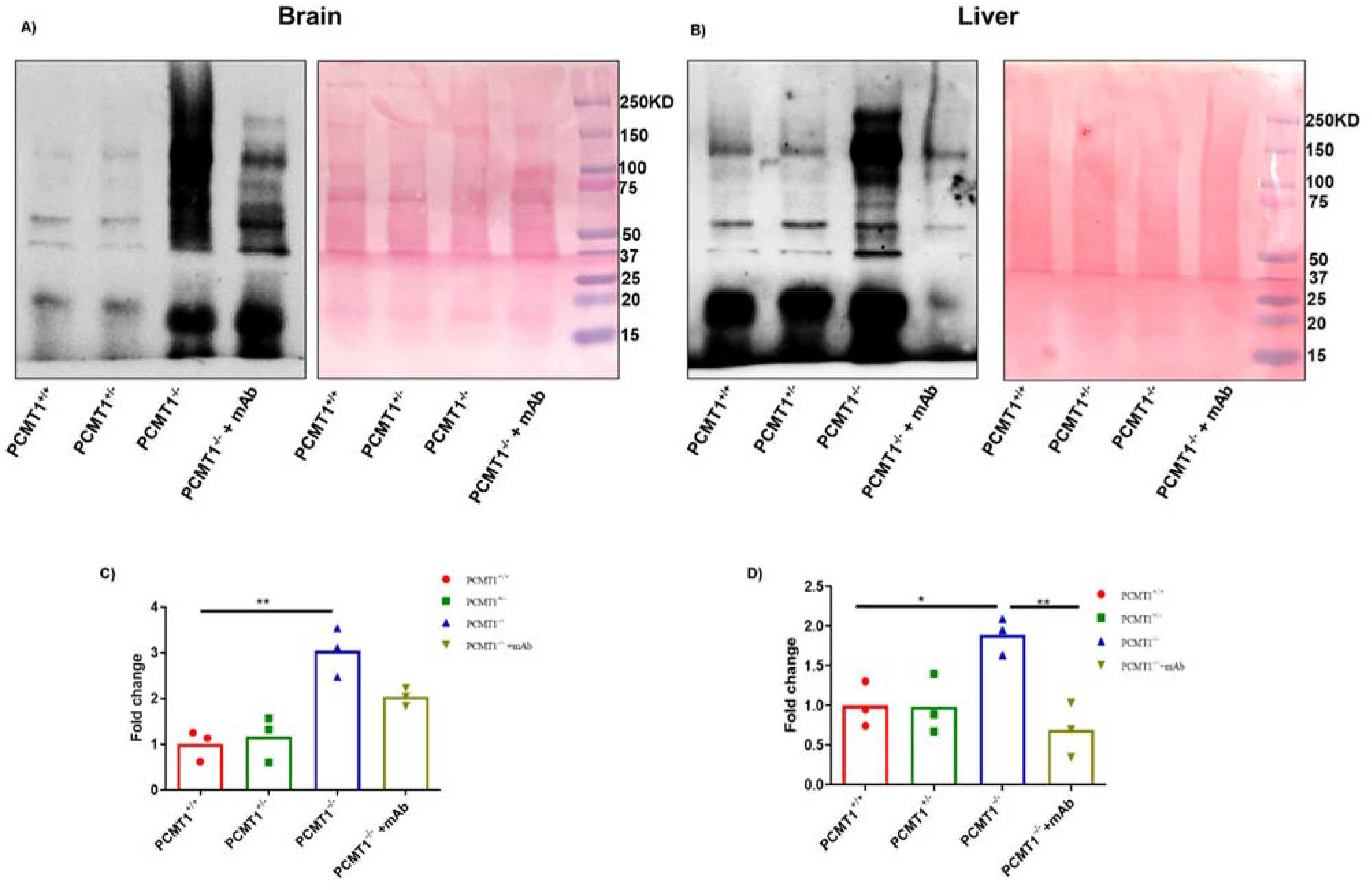
Anti-isoDGR mAb treatment reduces motif level in brain and liver from Pcmt1^-/-^ mice. Protein lysates from brain **(A)** and liver **(B)** of Pcmt1^+/+^, Pcmt1^+/-^, Pcmt1^-/-^, and mAb-treated Pcmt1^-/-^mice were subjected to western blot using isoDGR-specific antibody. Protein loadings were visualized by Ponceau S. Graphs show quantification of isoDGR-modified protein levels in mouse brain **(C)** and liver **(D).** Statistical significances were determined by 1-way ANOVA. ** p<0.01, * p<0.05.

### IsoDGR mAb reduces macrophage infiltration and inflammation of Pcmt1-KO liver

We previously identified that patients with atherosclerotic cardiovascular disease display marked isoDGR accumulation in ECM components of the vascular wall^13,23,24^, as well as in aortic tissues from 8-month-old PCMT1^+/+^ and Pcmt1^+/-^ mice. Immunofluorescent imaging revealed that isoDGR motifs positively correlated with CD68+ macrophages in the vessel wall,^13^ and this interaction was found to be mediated by integrins binding.^24^ Importantly, isoDGR-macrophage interactions triggered the secretion of several pro-inflammatory cytokines / chemokines including MCP1 and TNFα, thereby promoting vascular inflammation. Given our new finding that isoDGR-damaged proteins also significantly accumulate in the liver and brain of Pcmt1-KO mice, we next used immunofluorescent imaging to detect isoDGR-modified proteins and CD68+ macrophage distribution in the liver (Fig.3), spleen (Fig.S1), and thymus tissue (Fig.S2). Consistent with the western blot results, we observed a substantial accumulation of isoDGR in the liver, spleen, and thymus from Pcmt1^-/-^ mice that were significantly decreased by mAb treatment (Fig. 3A and 3B). In line with our hypothesis, isoDGR positively correlated with marked infiltration of CD68+ macrophages in each of the tissues analyzed (Fig. 3C). Moreover, the levels of CD68+ cells are proportionally increased with isoDGR accumulation in the tissues (Fig. 3D, S1D and S2D), suggesting that this motif promote macrophage recruitment to multiple organs including the liver, spleen, and thymus. Intriguingly, the lymphoid organs, as indicated by the slope values (spleen: slope=0.55, Fig.S1D; thymus: slope=0.20, Fig.S2D), are more sensitive to isoDGR-induced CD68+ cell infiltration than the liver (liver: slope=0.086, Fig.3D), suggesting that aging damaged isoDGR-proteins have a larger deleterious effect on these immune organs than the liver. The results imply that accumulation of isoDGR-proteins may contribute to inflammaging. However, Pcmt1^-/-^ mice treated with 1mg/kg/week anti-isoDGR mAb displayed a significant reduction in motif levels across all body tissues analyzed, with a concomitant decline in CD68+ macrophage infiltration (Fig. 3 A and C).

**Figure 3:**
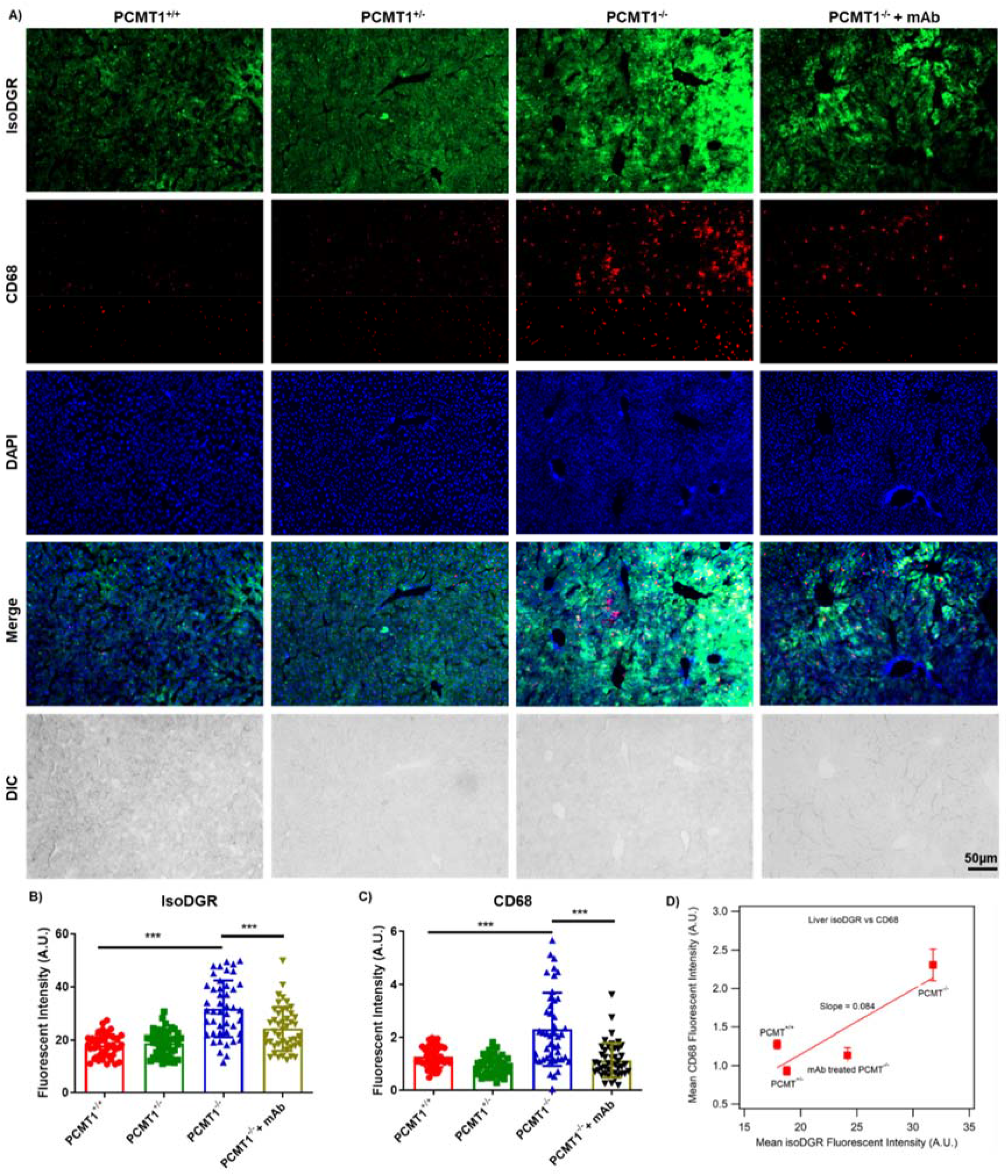
IsoDGR co-localization with CD68+ monocyte-macrophages in liver from Pcmt1-KO mice. **(A)** Representative immunostaining images showing isoDGR distribution and co-localisation with CD68+ macrophages in cryosectioned liver tissue (Pcmt1^+/+^, Pcmt1^+/-^ Pcmt1^-/-^, and mAb-treated Pcmt1^-/-^ mice at 6 weeks; n=3). **(B)** IsoDGR or **(C)** CD68 fluorescence in 50 randomized regions from 3 images of 3 independent spleen sections for each genotype were quantified using image J (graphs show averaged values for the same region from 3 images). **(D)**Plot showing the CD68 is proportionally increased with isoDGR accumulation, the slope indicates the sensitivity of the isoDGR-induced CD68+ cells infiltration. Statistical significances were determined by Kruskal-Wallis test. Results shown are mean values ± SE (***p<0.001).

### IsoDGR accumulation induces both local and systemic inflammation in Pcmt1^-/-^ mice

IsoDGR-damaged proteins are recognized by macrophage integrins and trigger pro-inflammatory cytokine release and cytotoxic functions, suggesting a potential role in the characteristic inflammaging in older individuals. Our previous findings also indicated that co-localization of isoDGR-modified proteins with CD68+ macrophages may be involved in chronic inflammation of the aorta in aged Pcmt1^+/-^ mice.^13^ As the elevated level of isoDGR-modified fibronectin can activate macrophages in Pcmt1^-/-^ to secrete proinflammatory cytokines/chemokines including MCP1 and TNFα, and recruit blood monocytes into tissues, we assessed the expression levels of inflammatory cytokines in tissues and organs of Pcmt1^-/-^ mice beyond known effects on the vasculature. To do this, we performed qRT-PCR using total RNA from the liver of 6-week-old Pcmt1^+/-^, Pcmt1^-/-^, mAb-treated Pcmt1^-/-^, and PCMT1^+/+^ mice. These results revealed significantly higher expression of mediators including MCP1, TNFα, and IL23 in the liver from Pcmt1^-/-^ mice, but mAb treatment effectively reduced this pro-inflammatory profile (Fig. 4A). To test whether the pro-inflammatory effects of isoDGR accumulation were local or systemic, we next assessed cytokine levels in blood plasma from Pcmt1^+/+^, Pcmt1^+/-^, Pcmt1^-/-^, and mAb-treated Pcmt1^-/-^ mice. Multiplex bead array assays confirmed that elevated concentrations of cytokines in plasma from Pcmt1^-/-^ mice could be reversed by anti-isoDGR immunotherapy. Individual mice with more extensive isoDGR accumulation (Fig. 2, 3, S1, and S2) also displayed corresponding higher levels of circulating pro-inflammatory cytokines (Fig 4).

**Figure 4:**
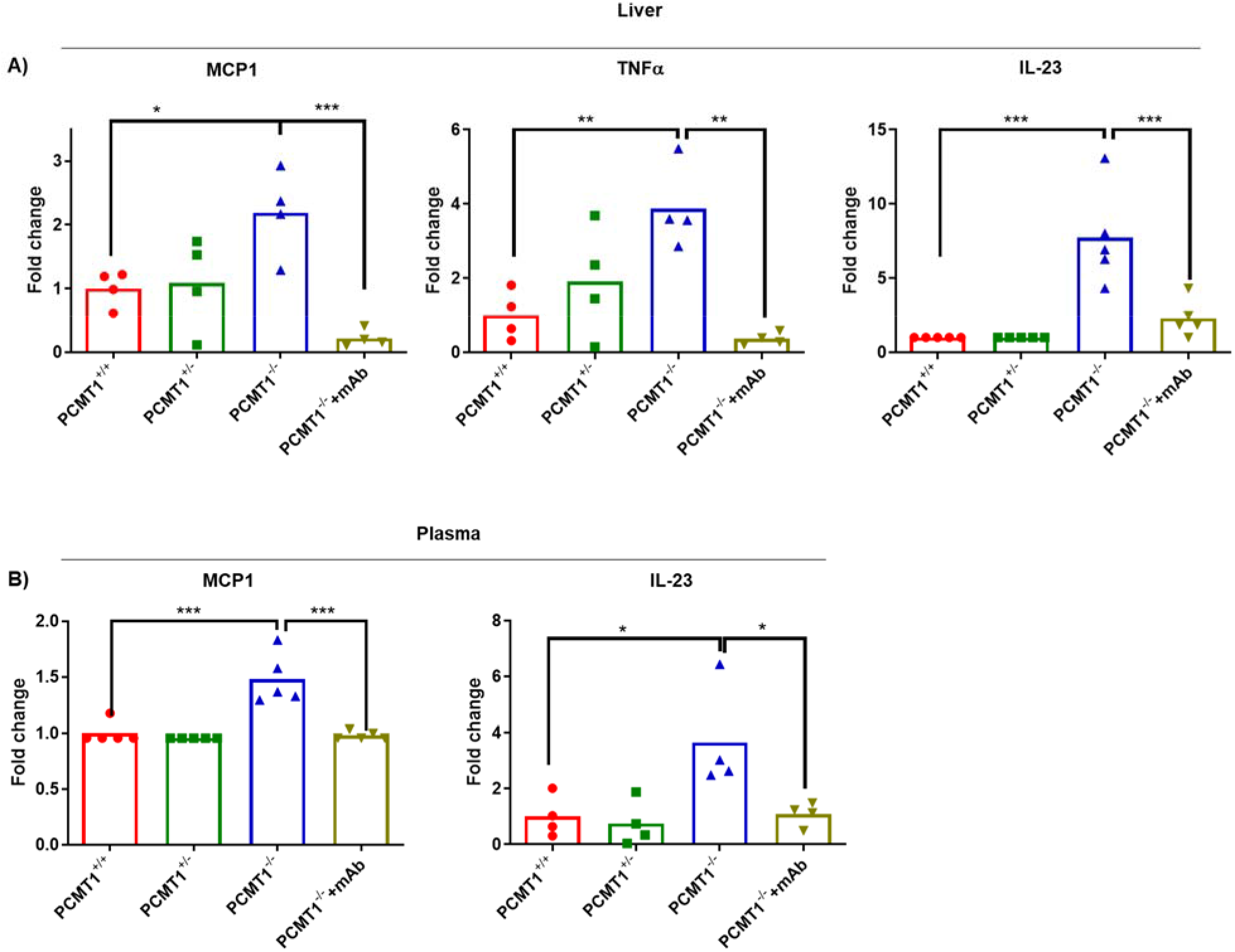
Elevated levels of local and systemic inflammation in Pcmt1^-/-^ mice. **(A)** Graph shows quantitative PCR analysis of pro-inflammatory cytokines in liver tissue from Pcmt1^+/+^, Pcmt1^+/-^, Pcmt1^-/-^, and mAb-treated Pcmt1^-/-^ mice at 6 weeks (n=4). **(B)** Graph shows cytokine quantification by multiplex bead array in blood plasma from Pcmt1^+/+^, Pcmt1^+/-^, Pcmt1^-/-^, and mAb-treated Pcmt1^-/-^ mice at 6 weeks (n=5). Statistical significances were determined by 1-way ANOVA, ***p<0.001, **p<0.01, *p<0.05.

### IsoDGR-modified proteins induce inflammation in young WT mice

To determine whether isoDGR can directly induce inflammation *in vivo*, we next injected C57BL/6 WT mice with isoDGR-modified plasma proteins (generated by incubating at pH 9 overnight to induce deamidation).^24^ After dialysis with 1X PBS, isoDGR-modified plasma proteins and unmodified WT control plasma were intravenously injected into 6-week-old WT mice. After 24h, we observed significantly higher levels of pro-inflammatory cytokines in the circulation of mice treated with isoDGR-plasma relative to WT plasma (Fig.S3A-F). Further analysis by immunostaining revealed that mice injected with deamidated plasma proteins also displayed increased levels of isoDGR-modified proteins in blood vessel walls that were positively correlated with CD68+ monocyte-macrophage infiltration (Fig.S3G). To further validate these findings, we next injected synthetic isoDGR peptide (Ac-GC(isoD)GRCGK) or PBS control into WT mice and collected blood plasma for assessment of pro-inflammatory cytokine levels 24h later. Again, we observed a significant elevation of plasma MCP1, TNFα, and IL1α concentrations in mice injected with isoDGR-peptide (Fig.S4). Together, these results suggest that isoDGR accumulation in blood plasma and body tissues can activate macrophages via integrin receptors to trigger the pathological release of pro-inflammatory cytokines and chemokines.

### Accumulation of isoDGR modified proteins in liver during ageing

We next sought to confirm that isoDGR can naturally accumulate in body tissues with advancing age due to the declining activity of the Pcmt1 enzyme. To do this, we used immunohistochemistry to interrogate isoDGR and CD68+ macrophage distribution in the liver from both PCMT1^+/+^ and Pcmt1^+/-^ mice at 4, 15, and 24 months (Fig.5A, B, C) (Pcmt1^-/-^ mice could not be assessed due to premature death of these animals). The results showed that isoDGR accumulates in the liver with advancing age, and to a greater extent in Pcmt1^+/-^ mice relative to PCMT1^+/+^ animals (Fig.5B, D). Pcmt1^+/-^ mice also displayed more extensive liver infiltration of CD68+ macrophages than PCMT1^+/+^ mice (Fig.5C, E), most likely due to more rapid decline in Pcmt1 function. Intriguingly, the accumulation of isoDGR-proteins is faster (steeper slope from 15-24 months, Fig.5D) in older mice (both PCMT1^+/+^ and Pcmt1^+/-^), suggesting the age-dependent risk of isoDGR induced pathology. Next, we analyzed inflammatory cytokine levels in plasma from 2-year-old PCMT1^+/+^ and Pcmt1^+/-^ mice. We detected significantly elevated concentrations of pro-inflammatory mediators in plasma from Pcmt1^+/-^ mice compared to PCMT1^+/+^ animals (Fig.S5). Together, these results indicate that increased levels of isoDGR-modified proteins with age promote cytokine release and likely contribute to the low-grade chronic inflammation that damages tissues and causes chronic diseases in the elderly.^35^

**Figure 5:**
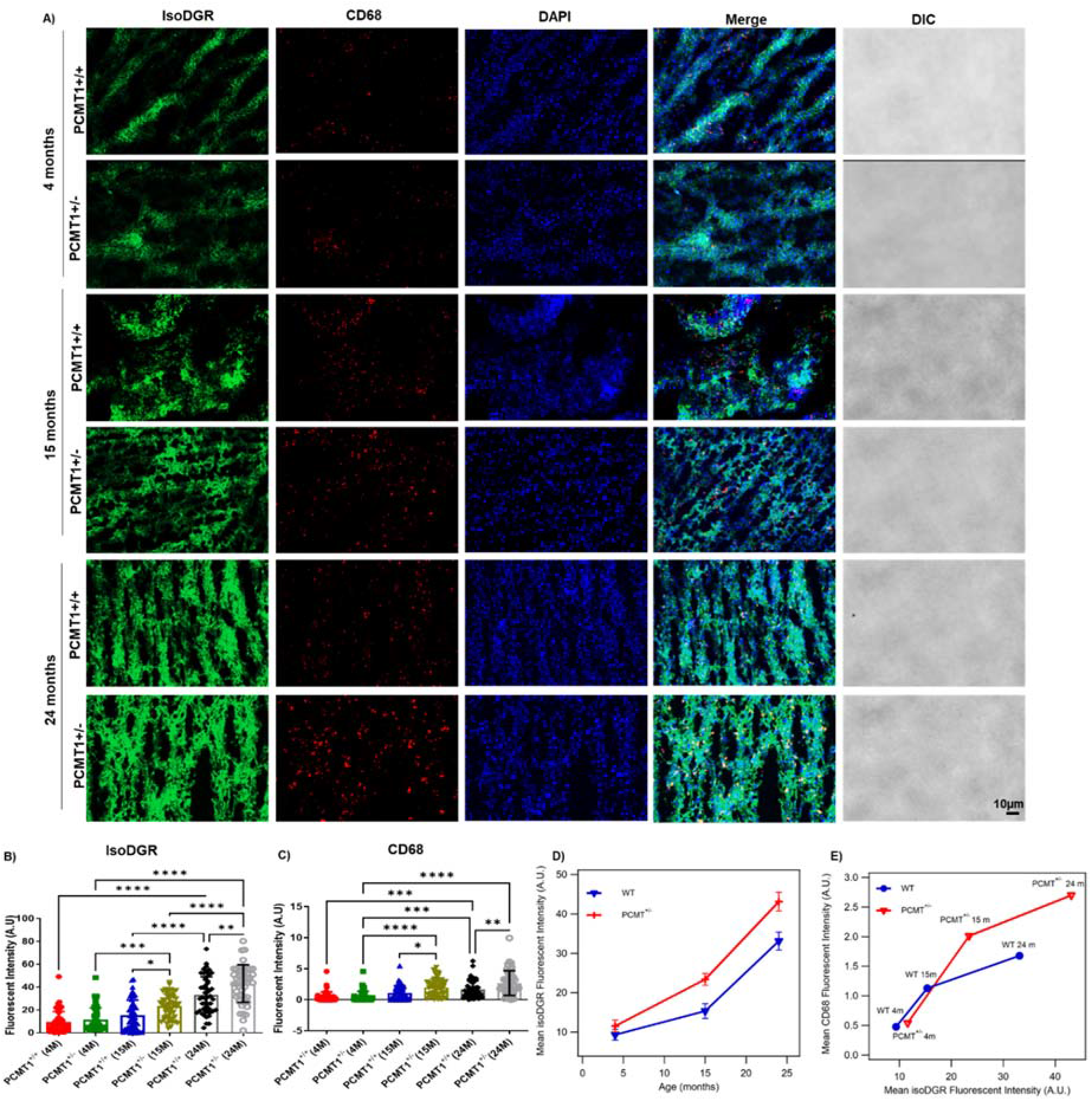
Age-induced accumulation of isoDGR co-localizes with CD68+ monocyte-macrophage infiltration of liver from PCMT1^+/+^ and Pcmt1^+/-^ mice. **(A)** Representative immunostaining of isoDGR protein distribution and co-localisation with CD68+ macrophages in cryosectioned liver tissue from Pcmt1^+/+^ and Pcmt1^+/-^ mice at 4, 15 and 24 months. IsoDGR **(B)** or CD68 **(C)** fluorescent intensities in a set of 50 random regions taken from 3 images representing 3 independent liver sections per genotype (quantified by image J software with average values plotted in the dot / bar graph). **(D)** Plot showing the accumulation of isoDGR-motif with age in both PCMT^+/+^ and PCMT^+/-^ animals. Results indicate that isoDGR accumulation is accelerated in older animal of both PCMT^+/+^ and PCMT^+/-^ mice. **(E)** CD68+ cells infiltration to tissues are proportional to the isoDGR levels. Statistical significances were determined by Kruskal-Wallis test. Results shown are mean values ± SE (**** p<0.0001, ***p<0.001, **p<0.01, *p<0.05).

### Immune clearance of isoDGR-damaged proteins

Weekly injection of isoDGR-specific mAb has significantly doubled the average lifespan of Pcmt1^-/-^ mice by reducing chronic inflammation. To understand the mechanistic underpinnings of the isoDGR-mAb mediated immunotherapy, we further tested how the mAb induced immune clearance of the damaged proteins by murine macrophage RAW cells using FACS and immunofluorescent imaging. RAW macrophages were seeded into 24-well plates and treated for 24h with 5μg/ml FITC-labelled isoDGR-fibronectin (isoDGR-FN-FITC), or native FN-FITC control, in the presence or absence of isoDGR-specific mAb (0 to 5μg/ml). After 45min incubation, excess isoDGR-FN-FITC was removed by washing with 1X PBS and quenched using trypan blue. FITC-positive phagocytic cells were then analyzed and quantified by FACS. This analysis revealed a clear mAb dose-dependent increase in phagocytic activity that was not observed with native FN-FITC control (data not shown) or in the absence of mAb (Fig. 6A), thus strongly indicating immune clearance of isoDGR-antigens by ADCP. Accumulation of isoDGR-damaged fibrinogen (isoDGR-FBG) has been identified in both atherosclerotic tissues and plasma from CVD patients,^13,24^ and we observed a similar mAb dose-dependent clearance when testing isoDGR-FBG antigen (Fig.6B).

**Figure 6:**
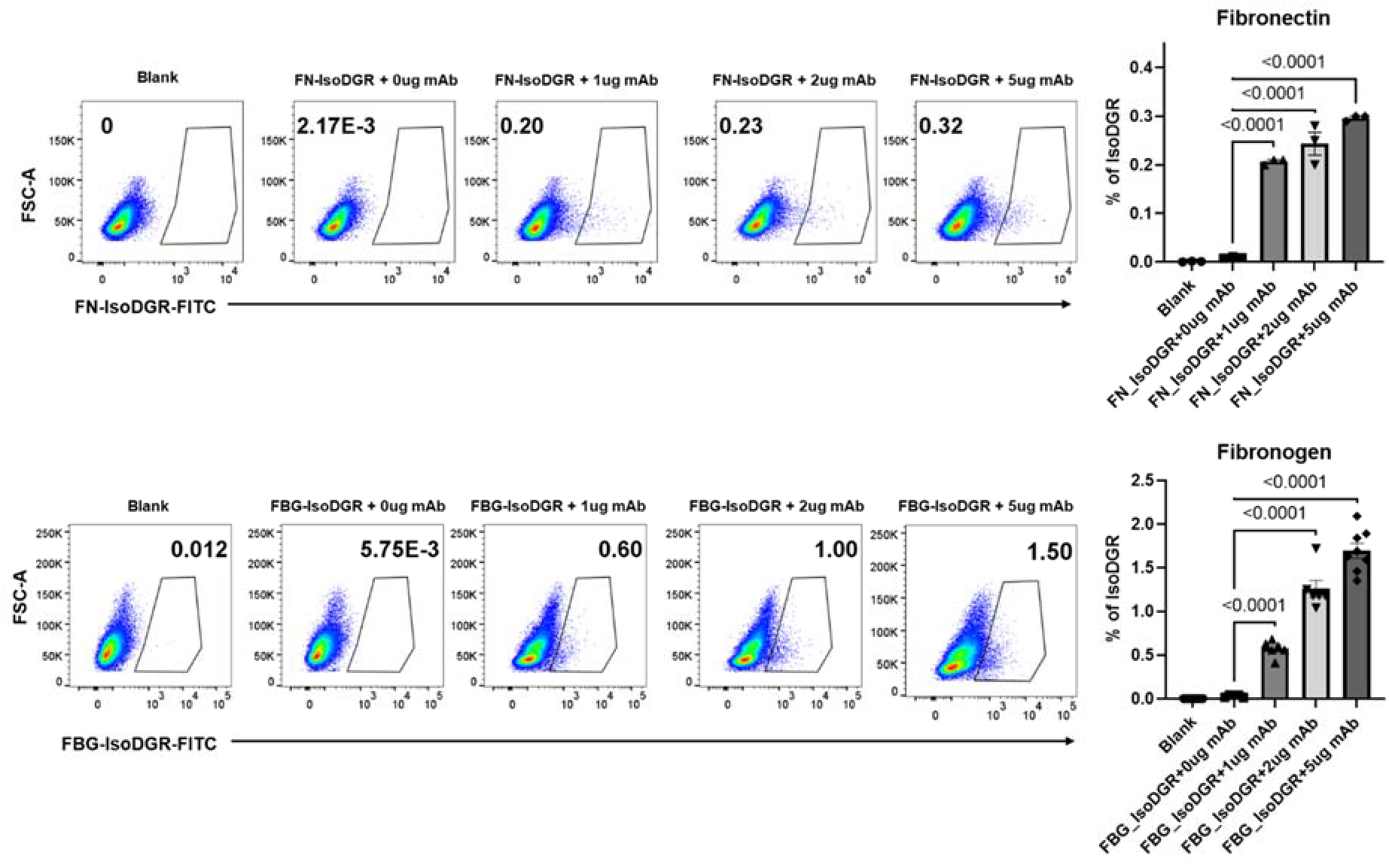
Motif-specific mAb promotes ADCP of isoDGR-modified fibronectin and fibrinogen. **(A)** Gating strategy used to quantify the number of phagocytic cells containing isoDGR-modified-fibronectin-FITC in the presence of varying concentration mAb treatment (1, 2 or 5μg/ml). Histogram and bar graph represent the average number of phagocytic RAW macrophages (n=3). **(B)** Gating strategy used to quantify the number of phagocytic cells containing isoDGR-modified-fibrinogen-FITC in the presence of varying concentration mAb treatment (1, 2 or 5μg/ml). Histogram and bar graph represent the average number of phagocytic RAW macrophages (n=7). Kruskal-Wallis test was used to assess statistical significances. Results shown are mean values ± SE (****p<0.0001, *** p<0.001, **p<0.01, *p<0.05).

These data indicate that the immune clearance observed was directed against isoDGR rather than being determined by protein identity. Indeed, we were also able to confirm mAb dose-dependent macrophage clearance of both isoDGR-FN and isoDGR-FBG using fluorescence microscopy (Fig.S6). Taken together, the data strongly suggest that target-specific mAb treatment stimulates macrophage clearance of isoDGR-modified proteins by ADCP and thereby reduces inflammatory cytokine expression in affected body tissues and extend the lifespan of the premature aging animals.

## DISCUSSION

Tissue aging is underpinned by progressive biomolecular damage that impairs the routine biochemical activities programmed by our genomes.^1,8,9^ Molecular damage to body tissues often results from spontaneous chemical processes, environmental stress, and the generation of reactive metabolites. In particular, non-enzymatic alterations to protein side chains have been strongly linked with the impairment of routine physiological processes.^2,5,7,18^ These degenerative protein modifications (DPMs) greatly increase the diversity of biomolecules/proteoforms present in body tissues, including the generation of species capable of mediating novel interactions, rather than simple loss of function.^24–26^ Among the different types of DPM, spontaneous protein deamidation has been proposed to represent ‘a molecular clock’ of tissue aging,^11^ but the underlying mechanisms have remained unclear and thus limited the development of interventions that target this DPM.

We previously developed custom proteomic techniques to enable a detailed study of global protein deamidation in complex biosamples.^28–30^ These techniques were used to investigate protein deamidation in vessel walls and plasma from patients with cardiovascular diseases (CVD) and cerebrovascular disease (CeVD). Intriguingly, we found that deamidation of extracellular matrix (ECM) proteins in these tissues confers paradoxical ‘gain-of-function’ changes by generating an isoDGR integrin binding sequence, thereby promoting leukocyte adhesion and pro-inflammatory cytokine expression^13,23,24^. Proteome-wide sequence analysis also showed that many ECM components featuring native ‘RGD’ integrin-binding motifs are highly enriched in NGR sequences susceptible to isoDGR modification. Thus, the risk of isoDGR-induced diseases will increase exponentially as protein repair functions decline with advancing age.

In the current study, we tested the concept that target specific immunotherapy can induce immune clearance of isoDGR-modified proteins *in vivo*, thus decreasing tissue inflammation and extending the lifespan of Pcmt1-KO mice (which lack the corresponding isoDGR repair enzyme). To this end, we generated hybridoma cells that produce isoDGR-specific mAb^13^ for use in immunoassays and to investigate the functional consequences of motif accumulation / antibody targeting in Pcmt1 KO mice. In this study, we observed a marked accumulation of isoDGR and elevated expression of pro-inflammatory cytokines in multiple body tissues from Pcmt1-KO mice. To confirm that isoDGR is responsible for these effects, we also induced deamidation of plasma proteins *in vitro* and showed that these could recapitulate pathology upon injection into young WT mice. Similar data were also obtained when injecting synthetic isoDGR-peptides into young WT mice. Conversely, a weekly injection of 1 mg/kg isoDGR-mAb was sufficient to induce immune clearance of isoDGR-modified proteins via ADCP. This not only doubled the lifespan of Pcmt1-KO mice but also preserved behaviour and coordinate functions and reduced pro-inflammatory cytokine levels in the circulation and body tissues. Together, these results indicate that mAb targeting of age-damaged proteins is a viable immunotherapeutic approach that could have wide-ranging implications for the treatment of chronic diseases prevalent in elderly populations.

Time-dependent biomolecular damage caused by DPMs (e.g. oxidation, deamidation, glycation, carbamylation) is well-established to mediate human aging and promote degenerative diseases. To date, the majority of research into combating age-related molecular damage has focused on the overexpression of specific repair enzymes (e.g. antioxidants) or restricting caloric intake to decrease levels of reactive metabolites. Our study clearly indicates that immunotherapeutic targeting of specific DPM triggers immune clearance of age-damaged proteins to reduce pathology and extend lifespan. We anticipate that future research in this area will uncover additional DPM-induced mechanisms of biological aging, as well as stimulate development of novel immune- and other therapies that target DPMs to increase human healthy lifespan.

## MATERIALS AND METHODS

### Animals

Mice deficient in deamidation repair enzyme Pcmt1 were previously generated by Clarke and co-worker^34^. We obtained Pcmt1^+/-^ (C57BL/6 background) mice from the Jackson Laboratory. Pcmt1^+/−^ (male and female) mice were bred, yielding litters comprising Pcmt1^+/+^, Pcmt1^+/−^, and Pcmt1^-/-^ offspring in the expected Mendelian ratios. Mouse genotype was confirmed by PCR using the following primers: oIMR1080:CGG CTG CAT ACG CTT GAT C, oIMR1081:CGA CAA GAC CGG CTT CCA T, oIMR1544:CAC GTG GGC TCC AGC ATT, oIMR3580:TCA CCA GTC ATT TCT GCC TTT G. Mice were maintained on normal chow diets and housed with regular light/dark cycles. All animal experiments used age- and sex-matched mice unless otherwise specified. All mouse procedures were performed in a humane manner and approved by the NTU Institutional Animal Care and Use Committee (IACUC protocol # ARF-SBS/NIE-A18016) or the Animal Care Committee at Brock University (AUP # 22-08-04).

### Anti-isoDGR immunotherapy

Pcmt1^-/-^ pups were intraperitoneally (i.p.) injected with 1mg/kg/week isoDGR-specific mAb ^13^ starting from 1 week old until use in the study. Age-matched control animals were given an equal volume of 1X PBS only. All mice were fed a normal laboratory diet (Altromin 1320M, Germany) from weaning onwards, body weight was measured weekly, and all animals were included in the survival analyses. Motor functional (hind-limb clasping test and ledge test) and behavioural functions were analyzed at 6 weeks. Blood was collected to prepare plasma for cytokine measurement.

### Hind-limb clasping test

Mice were suspended by their tails and the extent of hindlimb clasping was observed for 30 s. If both hind-limbs were splayed outward away from the abdomen with spread toes, a score of 0 was given. If one hind-limb was fully retracted or both hind-limbs partially retracted without touching the abdomen and with toes spread, a score of 1 was assigned. If both hind-limbs were partially retracted and in contact with the abdomen but without touching each other, a score of 2 was given. If both hind-limbs were fully clasped and touching the abdomen, a score of 3 was assigned.

### Ledge test

The ledge test for evaluating balance and coordination was carried out as previously described, with a minor modification. Mice were placed on a ledge (90 cm long, 0.5 cm wide, 20 cm high), and paw placement and forward movement were observed. The Time taken to cross the ledge and the number of paw slips were also recorded. Mice that left the ledge were excluded.

### IsoDGR-antigen uptake / ADCP assays

ADCP assays were performed by fluorescence microscopy and FACS. A total of 5 x 10^5^ murine RAW macrophages were seeded into 24-well plates and treated with 5 μg/ml FITC-labelled isoDGR-fibronectin (isoDGR-FN-FITC), FITC-labelled isoDGR fibrinogen (isoDGR-FG-FITC), or native FN-FITC / FG-FITC control for 24h.^24^ After 45 min incubation with varying doses of isoDGR-mAb (0 to 5 μg/ml), excess fluorescence was quenched by adding trypan blue and incubating for 10min. The cells were then washed 3 times and re-suspended in PBS. The mAb dose-dependent uptake of isoDGR-antigen by RAW cells was determined by FACS using FlowJo software to quantify phagocytic cells. To confirm the FACS results, cells were also examined by fluorescent imaging using a Zeiss LSM710 confocal microscope. The fluorescence intensity of each cell was calculated using Image J software.

### Histological assessment and immunostaining of aorta and liver tissues

Tissues from WT, Pcmt1^+/-^, Pcmt1^-/-^, and mAb-treated Pcmt1^-/-^ mice were collected and fixed with 4% PFA at 4 °C for 24 h. The tissues were washed with 1X PBS and transferred into 15% sucrose, followed by 30% sucrose, then stored at 4°C. Tissue was embedded in OCT compound with dry ice and cut into 10μm sections using A Leica CM3060S Cryostat. The sections were mounted onto Fisherbrand Superfrost plus microscope slides and kept in warm PBS for 20min to remove OCT prior to staining with haematoxylin and eosin. For immunostaining, slides were permeabilized with 0.5% PBST for 2-3h and then incubated with blocking buffer (2.5% normal goat serum, 1% BSA in 0.5% PBST) for 1h at RT prior to the addition of primary antibodies (isoDGR [1:200] and CD68 [1:200, Abcam, ab283654]) overnight at 4 °C. Slides were next washed with PBS (3X) for 5 min then incubated with secondary antibodies conjugated to AlexaFluor 488 and 594 (1:500) for 1h at RT. The slides were again washed with PBS (3X) for 5 min, then incubated with DAPI for 15 min to visualize cell nuclei. After staining, slides were washed with 1X PBS and mounted with aqueous mounting media. Images were acquired using a Zeiss LSM710 confocal microscope.

### Measurements of plasma cytokine levels

A cytokine multiplex bead assay, the LEGENDplex^™^ mouse inflammation panel (Biolegend, San Diego, CA), was used to measure 13 different cytokines (IL23, IL1α, IL1β, IL6, IL10, IL12p70, IL17A, IL23, IL27, MCP1, IFNβ, IFNγ, TNFα, and GMCSF) in blood plasma from WT, Pcmt1^+/-^, Pcmt1^-/-^ and mAb-treated Pcmt1^-/-^ animals (assessed by LSRII flow cytometer according to the manufacturer’s protocol).

### Real-time PCR

Total RNA isolated from tissues was treated with DNase and reverse-transcribed using a first-strand DNA synthesis kit from Invitrogen. The PCR was performed on an ABI Fast 7500 System (Applied Biosystems, Foster City, CA). TaqMan probes for the respective genes were custom-generated by Applied Biosystems based on the sequences in the Illumina array and used as per the manufacturer’s instructions. Expression levels of target genes were determined in triplicate from the standard curve and normalized to GAPDH mRNA level.

### Western blot analysis

Western blot analysis was performed using standard methods. Primary antibodies and dilutions were as follows: isoDGR (mouse monoclonal 1:1000), Pcmt1 (rabbit polyclonal, Abcam 1:1000), GAPDH (Invitrogen 1:1000).

### Statistics

Statistical analyses were performed using GraphPad Prism v9.0 (GraphPad Software, Inc., San Diego, CA). Statistical differences in Kaplan-Meier survival curves between experimental groups were evaluated using Mantel-Cox log-rank test. Data were tested for normality using D’Agostino & Pearson, or Sapiro-Wilk tests. For normally distributed data, differences between groups were assessed either by two-tailed unpaired Student’s t-test or one-way ANOVA for multiple group analyses, followed by Tukey’s multiple comparisons test. Otherwise, Kruskal-Wallis test with Dunn’s multiple comparisons post hoc test was used.*p* < 0.05 was considered significant.

## AUTHOR CONTRIBUTIONS

K.P., N.M. and S.K.S. designed the project. K.P., K.M., S.C.N., R.I., E.M., T,L., G.W.Z., B.S., and T.G. performed the experiments and data analysis; P.K. drafted the manuscript; A.M., R.E.K.M., E.L.T., D.D.O., Y.G.G., K.L.L., I.H.S. A.M.R, R.K. and C.C. provided resources/equipment and revised the manuscript; Y.G.G., K.L.L. and S.K.S contributed grants/reagents/materials/analysis tools. S.K.S. conceived and supervised the project. N.M. and S.K.S. revised the manuscript.

## DATA AVAILABILITY

All data presented in this study are included in the main text or the supplemental materials.

## REFERENCES

1 Hipp, M. S., Kasturi, P. & Hartl, F. U. The proteostasis network and its decline in ageing. Nature reviews. Molecular cell biology 20, 421–435, doi:10.1038/s41580-019-0101-y (2019).

2 Gallart-Palau, X. et al. Degenerative protein modifications in the aging vasculature and central nervous system: A problem shared is not always halved. Ageing research reviews 53, 100909, doi:10.1016/j.arr.2019.100909 (2019).

3 Maksimovic, I. & David, Y. Non-enzymatic Covalent Modifications as a New Chapter in the Histone Code. Trends in biochemical sciences, doi:10.1016/j.tibs.2021.04.004 (2021).

4 Delanghe, S., Delanghe, J. R., Speeckaert, R., Van Biesen, W. & Speeckaert, M. M. Mechanisms and consequences of carbamoylation. Nature reviews. Nephrology 13, 580–593, doi:10.1038/nrneph.2017.103 (2017).

5 Gallart-Palau, X., Serra, A. & Sze, S. K. Uncovering Neurodegenerative Protein Modifications via Proteomic Profiling. Int Rev Neurobiol 121, 87–116, doi:10.1016/bs.irn.2015.06.002 (2015).

6 Dai, Z., Ramesh, V. & Locasale, J. W. The evolving metabolic landscape of chromatin biology and epigenetics. Nature reviews. Genetics 21, 737–753, doi:10.1038/s41576-020-0270-8 (2020).

7 Truscott, R. J. W., Schey, K. L. & Friedrich, M. G. Old Proteins in Man: A Field in its Infancy. Trends in biochemical sciences 41, 654–664, doi:10.1016/j.tibs.2016.06.004 (2016).

8 Clarke, S. Aging as war between chemical and biochemical processes: protein methylation and the recognition of age-damaged proteins for repair. Ageing research reviews 2, 263–285, doi:10.1016/s1568-1637(03)00011-4 (2003).

9 da Costa, J. P. et al. A synopsis on aging-Theories, mechanisms and future prospects. Ageing research reviews 29, 90–112, doi:10.1016/j.arr.2016.06.005 (2016).

10 Gladyshev, V. N. et al. Molecular damage in aging. Nature Aging 1, 1096–1106, doi:10.1038/s43587-021-00150-3 (2021).

11 Robinson, N. E. & Robinson, A. B. Molecular clocks. Proceedings of the National Academy of Sciences of the United States of America 98, 944–949, doi:10.1073/pnas.98.3.944 (2001).

12 McKerrow, J. H. & Robinson, A. B. Primary sequence dependence of the deamidation of rabbit muscle aldolase. Science (New York, N.Y.) 183, 85, doi:10.1126/science.183.4120.85 (1974).

13 Park, J. E. et al. Aging-induced isoDGR-modified fibronectin activates monocytic and endothelial cells to promote atherosclerosis. Atherosclerosis 324, 58–68, doi:10.1016/j.atherosclerosis.2021.03.020 (2021).

14 Kim, E., Lowenson, J. D., MacLaren, D. C., Clarke, S. & Young, S. G. Deficiency of a protein-repair enzyme results in the accumulation of altered proteins, retardation of growth, and fatal seizures in mice. Proceedings of the National Academy of Sciences of the United States of America 94, 6132–6137, doi:10.1073/pnas.94.12.6132 (1997).

15 Geiger, T. & Clarke, S. Deamidation, isomerization, and racemization at asparaginyl and aspartyl residues in peptides. Succinimide-linked reactions that contribute to protein degradation. The Journal of biological chemistry 262, 785–794 (1987).

16 Yamamoto, A. et al. Deficiency in protein L-isoaspartyl methyltransferase results in a fatal progressive epilepsy. The Journal of neuroscience: the official journal of the Society for Neuroscience 18, 2063–2074, doi:10.1523/jneurosci.18-06-02063.1998 (1998).

17 Adav, S. S. et al. iTRAQ quantitative clinical proteomics revealed role of Na(+)K(+)-ATPase and its correlation with deamidation in vascular dementia. J Proteome Res 13, 4635–4646, doi:10.1021/pr500754j (2014).

18 Adav, S. S. & Sze, S. K. Insight of brain degenerative protein modifications in the pathology of neurodegeneration and dementia by proteomic profiling. Molecular Brain 9, 92 (2016).

19 Kim, J. et al. New findings on SNP variants of human protein L-isoaspartyl methyltransferase that affect catalytic activity, thermal stability, and aggregation. PloS one 13, e0198266, doi:10.1371/journal.pone.0198266 (2018).

20 Juang, C. et al. Polymorphic Variants of Human Protein l-Isoaspartyl Methyltransferase Affect Catalytic Activity, Aggregation, and Thermal Stability: IMPLICATIONS FOR THE ETIOLOGY OF NEUROLOGICAL DISORDERS AND COGNITIVE AGING. The Journal of biological chemistry 292, 3656–3665, doi:10.1074/jbc.M116.765222 (2017).

21 Qin, Z., Dimitrijevic, A. & Aswad, D. W. Accelerated protein damage in brains of PIMT+/-mice; a possible model for the variability of cognitive decline in human aging. Neurobiology of aging 36, 1029–1036, doi:10.1016/j.neurobiolaging.2014.10.036 (2015).

22 Hao, P., Adav, S. S., Gallart-Palau, X. & Sze, S. K. Recent advances in mass spectrometric analysis of protein deamidation. Mass spectrometry reviews 36, 677–692, doi:10.1002/mas.21491 (2017).

23 Cheow, E. S. H. et al. The Role of Protein Deamidation in Cardiovascular Disease. Proceedings of the 23rd American Peptide Symposium 17, 212–213 (2013).

24 Dutta, B. et al. Monocyte adhesion to atherosclerotic matrix proteins is enhanced by Asn-Gly-Arg deamidation. Scientific Reports 7, 5765, doi:10.1038/s41598-017-06202-2 (2017).

25 Curnis, F. et al. Spontaneous Formation of L-Isoaspartate and Gain of Function in Fibronectin. Journal of Biological Chemistry 281, 36466–36476, doi:10.1074/jbc.M604812200 (2006).

26 Corti, A. & Curnis, F. Isoaspartate-dependent molecular switches for integrin-ligand recognition. Journal of cell science 124, 515–522, doi:10.1242/jcs.077172 (2011).

27 Spitaleri, A. et al. Structural basis for the interaction of isoDGR with the RGD-binding site of alphavbeta3 integrin. The Journal of biological chemistry 283, 19757–19768, doi:10.1074/jbc.M710273200 (2008).

28 Hao, P., Ren, Y., Alpert, A. J. & Sze, S. K. Detection, evaluation and minimization of nonenzymatic deamidation in proteomic sample preparation. Molecular & cellular proteomics: MCP 10, O111.009381, doi:10.1074/mcp.O111.009381 (2011).

29 Sze, S. K., JebaMercy, G. & Ngan, S. C. Profiling the ‘deamidome’ of complex biosamples using mixed-mode chromatography-coupled tandem mass spectrometry. Methods (San Diego, Calif.), doi:10.1016/j.ymeth.2020.05.005 (2020).

30 Hao, P., Adav, S. S., Gallart-Palau, X. & Sze, S. K. Recent advances in mass spectrometric analysis of protein deamidation. Mass spectrometry reviews 36, 677–692, doi:10.1002/mas.21491 (2017).

31 Robinson, N. E. & Robinson, A. B. Deamidation of human proteins. Proceedings of the National Academy of Sciences of the United States of America 98, 12409–12413, doi:10.1073/pnas.221463198 (2001).

32 Smith, L. M. et al. A five-level classification system for proteoform identifications. Nature methods 16, 939–940, doi:10.1038/s41592-019-0573-x (2019).

33 Hao, P. et al. Deep proteomic profiling of human carotid atherosclerotic plaques using multidimensional LC-MS/MS. Proteomics. Clinical applications 8, 631–635, doi:10.1002/prca.201400007 (2014).

34 Kim, E., Lowenson, J. D., Clarke, S. & Young, S. G. Phenotypic analysis of seizure-prone mice lacking L-isoaspartate (D-aspartate) O-methyltransferase. The Journal of biological chemistry 274, 20671–20678, doi:10.1074/jbc.274.29.20671 (1999).

35 Franceschi, C., Garagnani, P., Parini, P., Giuliani, C. & Santoro, A. Inflammaging: a new immune-metabolic viewpoint for age-related diseases. Nature reviews. Endocrinology 14, 576–590, doi:10.1038/s41574-018-0059-4 (2018).

36 Liberale, L., Montecucco, F., Tardif, J. C., Libby, P. & Camici, G. G. Inflamm-ageing: the role of inflammation in age-dependent cardiovascular disease. European heart journal 41, 2974–2982, doi:10.1093/eurheartj/ehz961 (2020).

37 Ferrucci, L. & Fabbri, E. Inflammageing: chronic inflammation in ageing, cardiovascular disease, and frailty. Nature reviews. Cardiology 15, 505–522, doi:10.1038/s41569-018-0064-2 (2018).

38 Nimmerjahn, F., Gordan, S. & Lux, A. FcγR dependent mechanisms of cytotoxic, agonistic, and neutralizing antibody activities. Trends in immunology 36, 325–336, doi:10.1016/j.it.2015.04.005 (2015).

39 Lowenson, J. D., Kim, E., Young, S. G. & Clarke, S. Limited accumulation of damaged proteins in l-isoaspartyl (D-aspartyl) O-methyltransferase-deficient mice. The Journal of biological chemistry 276, 20695–20702, doi:10.1074/jbc.M100987200 (2001).

40 Cahill, L. S. et al. Aged hind-limb clasping experimental autoimmune encephalomyelitis models aspects of the neurodegenerative process seen in multiple sclerosis. Proceedings of the National Academy of Sciences of the United States of America 116, 22710–22720, doi:10.1073/pnas.1915141116 (2019).

41 Castillo-Mariqueo, L. & Giménez-Llort, L. Clasping, ledge-score coordination and early gait impairments as primary behavioural markers of functional impairment in Alzheimer’s disease. Behavioural brain research 435, 114054, doi:10.1016/j.bbr.2022.114054 (2022).

42 Schoonover, K. E. et al. Interactions between knockout of schizophrenia risk factor Dysbindin-1 and copper metabolism in mice. Brain research bulletin 164, 339–349, doi:10.1016/j.brainresbull.2020.07.024 (2020).

43 Crowe, J. E., Jr. Human Antibodies for Viral Infections. Annual review of immunology 40, 349–386, doi:10.1146/annurev-immunol-042718-041309 (2022).

44 Pinto, S., Pahl, J., Schottelius, A., Carter, P. J. & Koch, J. Reimagining antibody-dependent cellular cytotoxicity in cancer: the potential of natural killer cell engagers. Trends in immunology 43, 932–946, doi:10.1016/j.it.2022.09.007 (2022).

45 Biburger, M., Lux, A. & Nimmerjahn, F. How immunoglobulin G antibodies kill target cells: revisiting an old paradigm. Advances in immunology 124, 67–94, doi:10.1016/b978-0-12-800147-9.00003-0 (2014).

