## Supplemental Figures for "Antibody targeting of aging damaged isoDGR-proteins doubles lifespan in a mouse model of chronic inflammation"

### Supplementary Figures

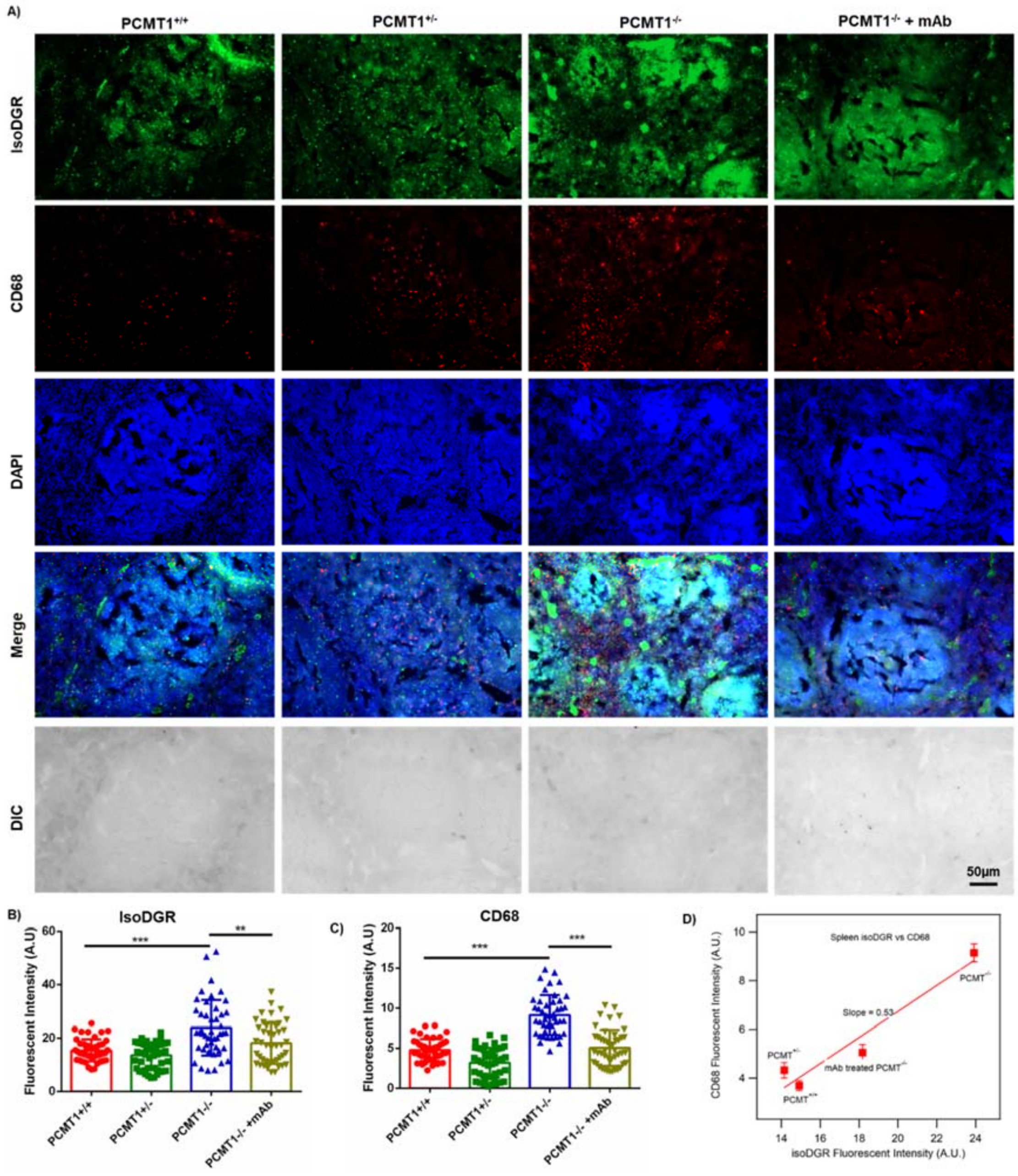


***Figure S1: Increased isoDGR levels/co-localization with CD68+ macrophages in Pcmt1$^{-/-}$ mouse spleen (A)*** *Representative immunostaining images showing isoDGR distribution and co-localisation with CD68+ macrophages in cryosectioned spleen tissue (Pcmt1+/+, Pcmt1+/-, Pcmt1-/-, and mAb-treated Pcmt1-/- mice at 6 weeks).* ***(B)*** *IsoDGR or* ***(C)*** *CD68 fluorescence in 50 randomized regions from 3 images of 3 independent spleen sections for each genotype were quantified using image J (graphs shows averaged values for the same region from 3 images).* ***(D)****Plot showing the CD68 is proportionally increased with isoDGR accumulation with slope=0.55, suggesting spleen is highly sensitive to aging-damaged isoDGR-molecules. One-way ANOVA was used to assess statistical significances. Results shown are mean values ± SE (*** p<0.001, ** p<0.01).*

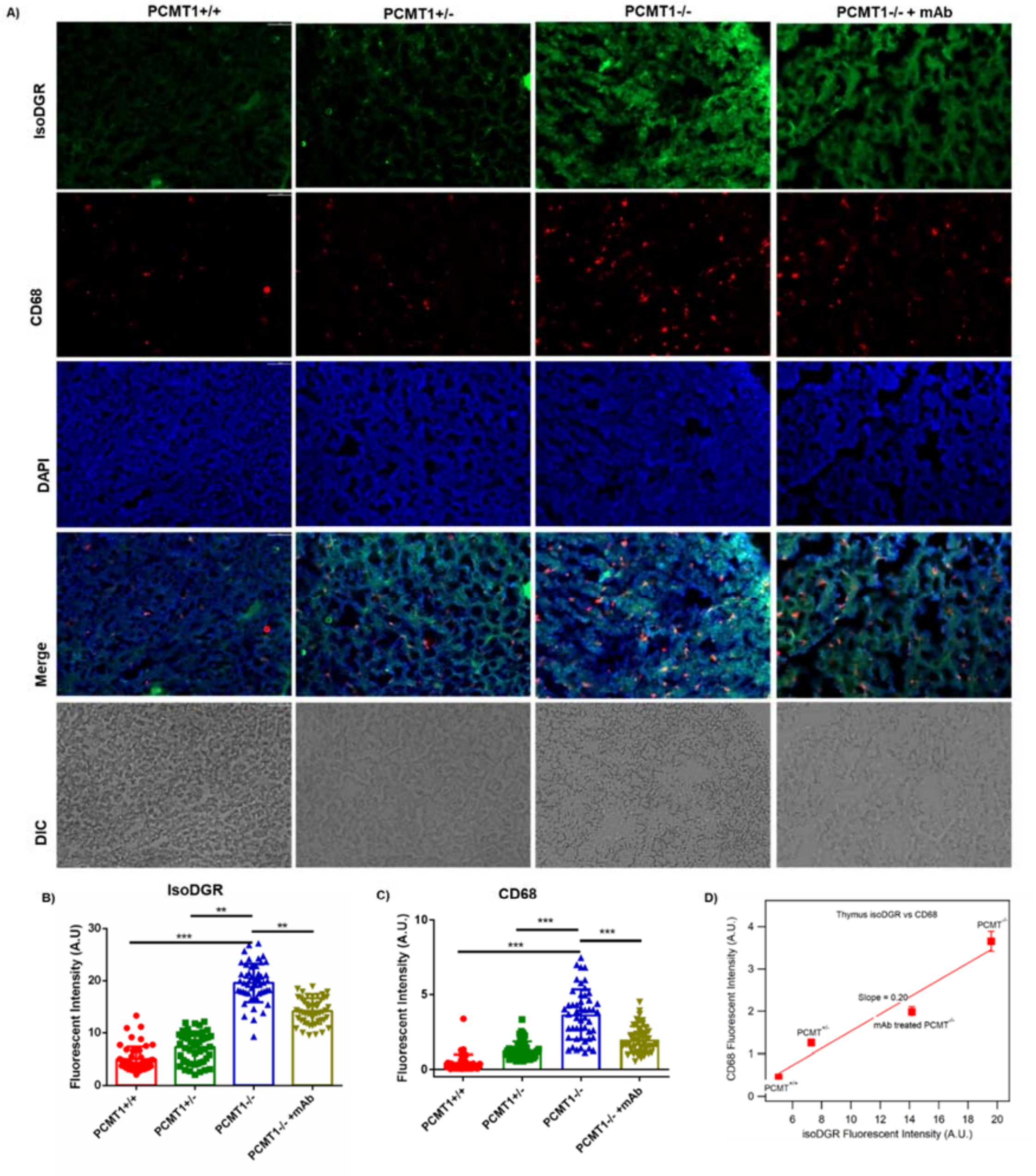


***Figure S2: Increased isoDGR levels/co-localization with CD68+ macrophages in Pcmt1$^{-/-}$ mouse thymus* (A)** *Representative immunostaining images showing isoDGR distribution and co-localisation with CD68+ macrophages in cryosectioned thymus tissue (Pcmt1$^{+/+}$, Pcmt1$^{+/-}$, Pcmt1$^{-/-}$, and mAb-treated Pcmt1$^{-/-}$ mice at 6 weeks).* ***(B)*** *IsoDGR or* ***(C)*** *CD68 fluorescence in 50 randomized regions from 3 images of 3 independent thymus sections for each genotype were quantified using image J (graphs shows averaged values for the same region from 3 images).* ***(D)****Plot showing the CD68 is proportionally increased with isoDGR accumulation with slope=0.20. Kruskal-Wallis test was used to calculate statistical significances. Results shown are mean values ± SE (*** $p<0.001$, ** $p<0.01$).*

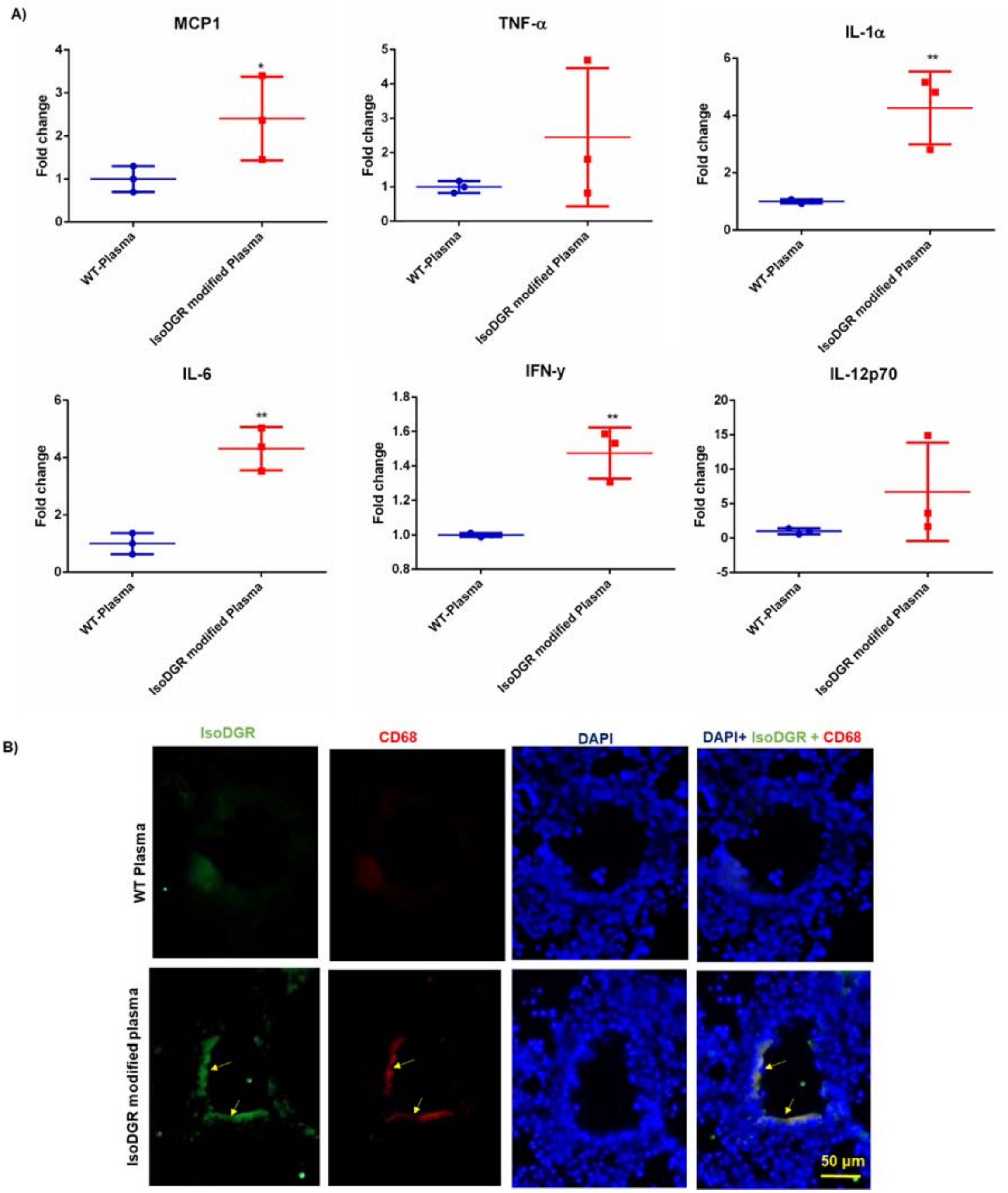


***Figure S3: IsoDGR-modified plasma proteins induce systemic inflammation in wild-type C57BL/6 mice (A)*** *Graph shows quantitation of blood plasma cytokines in WT mice injected with unmodified plasma or isoDGR-modified plasma (n=3).* ***(B)*** *Representative immunostaining images showing isoDGR protein distribution and co-localisation with CD68+ macrophages in cryosectioned lung small vessels from WT mice treated with WT or isoDGR-modified plasma. Statistical significances were assessed by one-way ANOVA. Results shown are values mean ± SE (** $p<0.01$, * $p<0.05$).*

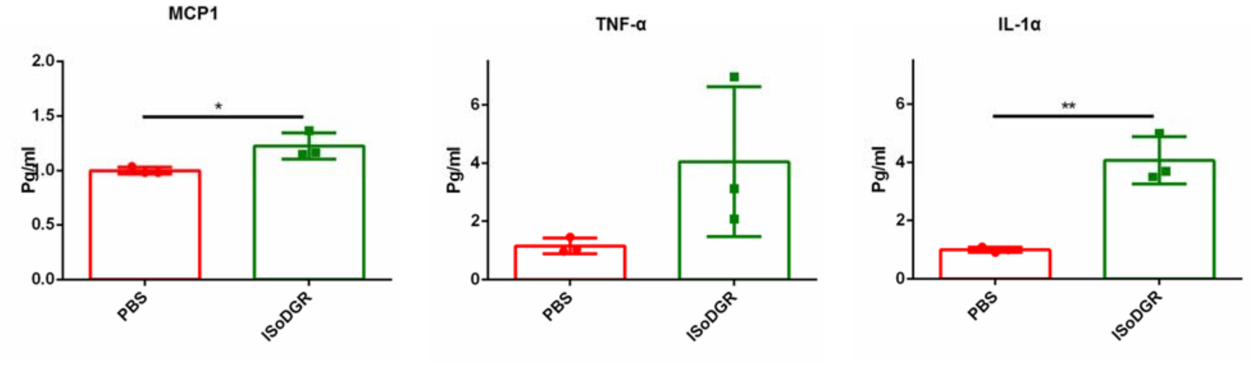


***Figure S4: IsoDGR-peptides promote inflammatory cytokine release in wild-type C57BL/6 mice***
*Graph shows quantitation of blood plasma chemokines / cytokines MCP1* ***(A)*** *TNF-α* ***(B)*** *and IL1α* ***(C)*** *in C57BL6 mice treated with either isoDGR synthetic peptide or PBS-only control (n=3). Statistical significances were calculated by one-way ANOVA. Results shown are mean values ± SE (** $p<0.01$, * $p<0.05$).*

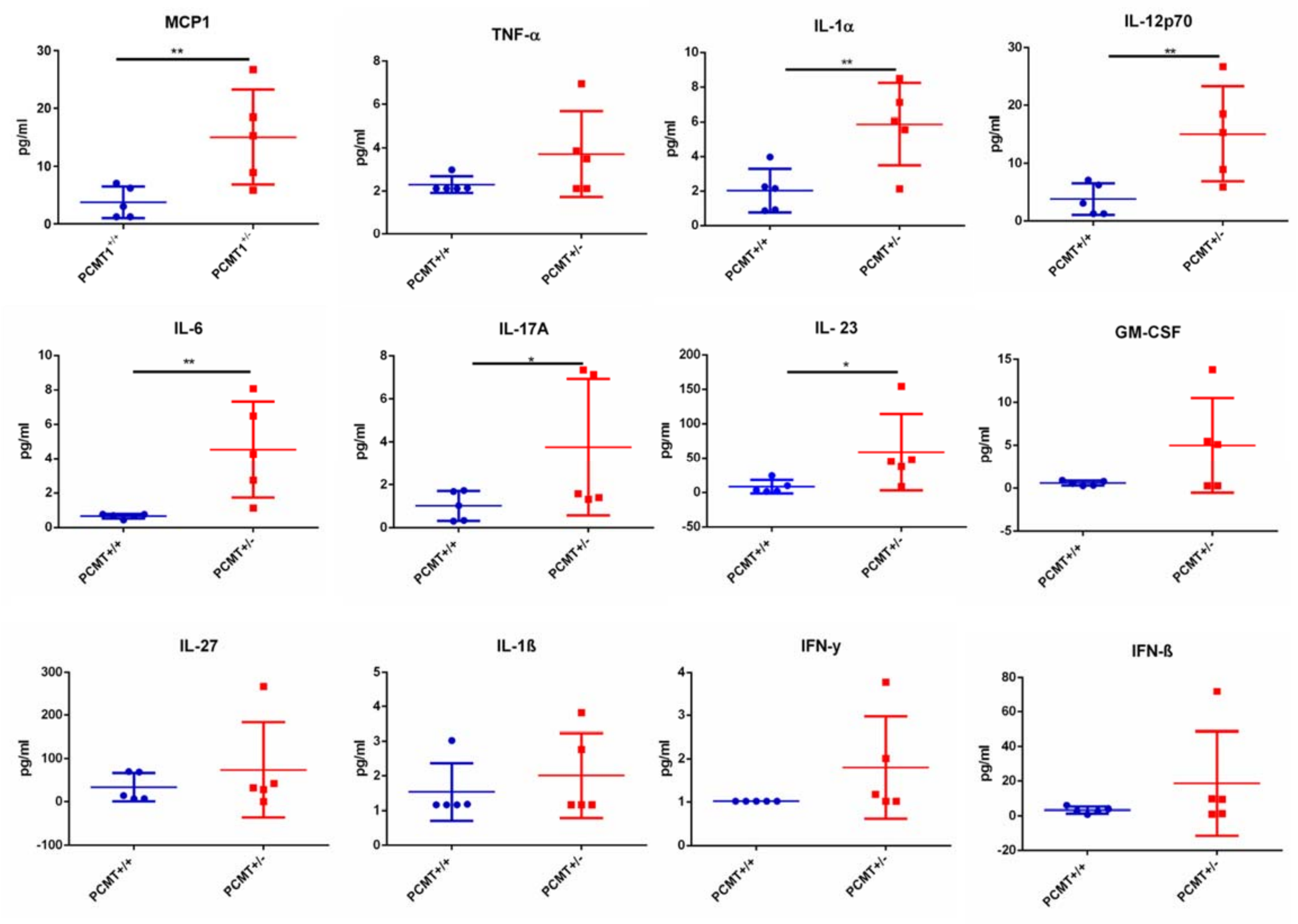


***Figure S5: Partial deletion of Pcmt1 leads to systemic elevation of proinflammatory cytokines in old mice***

*Graph shows quantification of inflammatory cytokines in plasma from 2-year-old Pcmt1$^{+/+}$ and Pcmt1$^{+/-}$ mice (n=5). Statistical significances were assessed by one-way ANOVA. Results shown are mean values ± SE (** p<0.01, * p<0.05).*

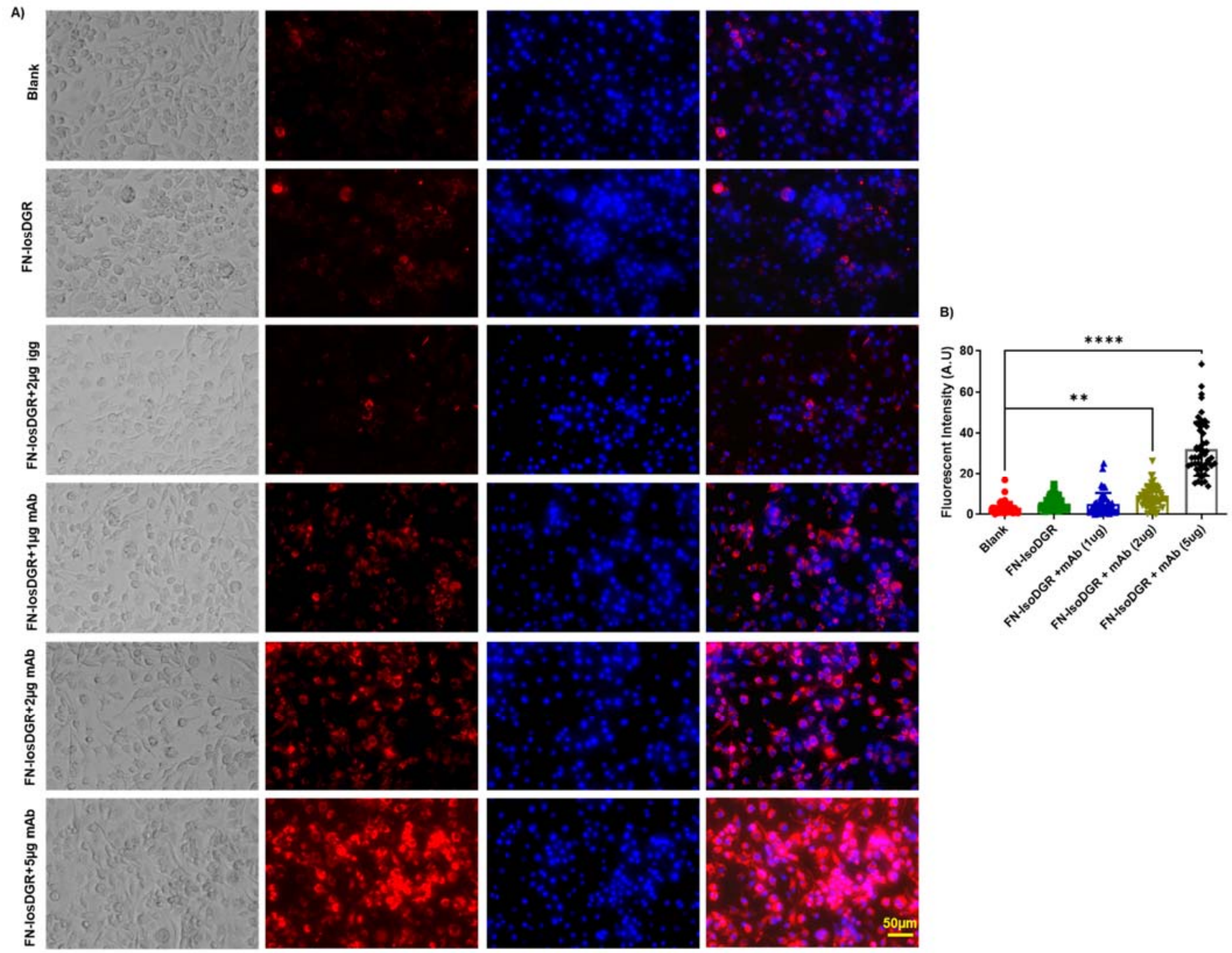


***Figure S6: Target-specific mAb enhances phagocytosis of isoDGR-modified fibronectin.*** *Representative immunostaining images showing that phagocytosis of isoDGR-FN by RAW macrophages increases with motif-specific mAb concentration (1-5µg/ml). Bar graph shows average number of phagocytic RAW macrophages. Fluorescent signal in 50 random regions taken from 3 images in 3 independent experiments were quantified by image J software (graph shows averaged values). Kruskal-Wallis test was used to calculate statistical significances. Results shown are mean values ± SE (*** $p<0.001$, ** $p<0.01$).*

Full WB images of Fig. 1A

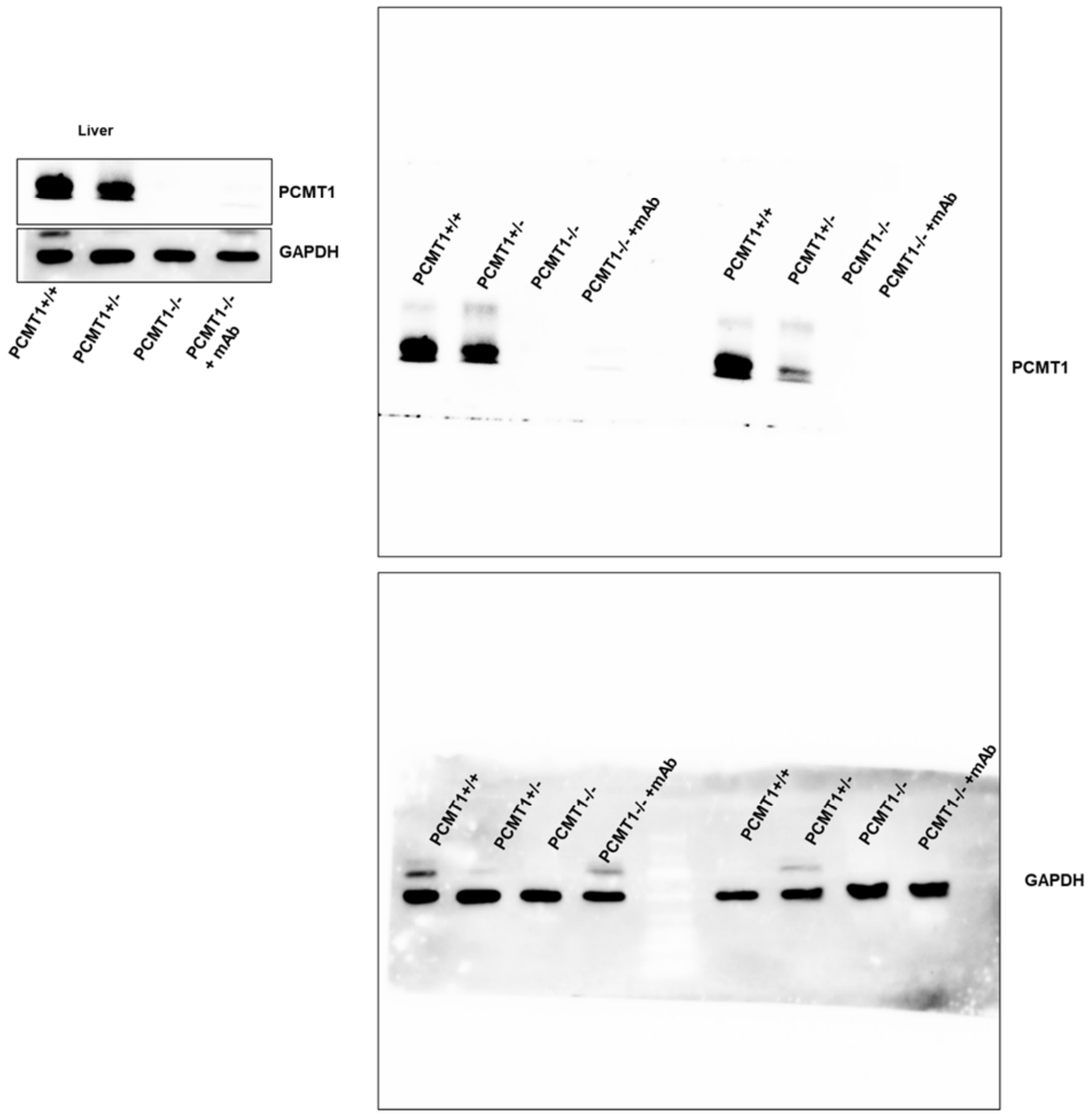
